# Maternal age affects equine Day 8 embryo gene expression both in trophoblast and inner cell mass

**DOI:** 10.1101/2021.04.07.438786

**Authors:** E. Derisoud, L. Jouneau, C. Dubois, C. Archilla, Y. Jaszczyszyn, R. Legendre, N. Daniel, N. Peynot, M. Dahirel, J. Auclair-Ronzaud, L. Wimel, V. Duranthon, P. Chavatte-Palmer

**Affiliations:** Université Paris-Saclay, UVSQ, INRAE, BREED, 78350, Jouy-en-Josas, France; Ecole Nationale Vétérinaire d’Alfort, BREED, 94700, Maisons-Alfort, France; IFCE, Plateau technique de la Valade, Chamberet, France; Institute for Integrative Biology of the Cell (I2BC), UMR 9198 CNRS, CEA, Paris-Sud University F-91198, Gif-sur-Yvette, France; Institut Pasteur—Bioinformatics and Biostatistics Hub—Department of Computational Biology, USR 3756 IP CNRS, Paris, France

**Author notes:** Corresponding author **Correspondence:** Emilie Derisoud, Domaine de Vilvert, Bâtiment 230-231, UMR 1198 BREED, INRAE, 78350, Jouy-en-Josas, FRANCE, Pascale Chavatte-Palmer, Domaine de Vilvert, Bâtiment 230-231, UMR 1198 BREED, INRAE, 78350, Jouy-en-Josas, France.

## Abstract

**Background:** Breeding a mare until she is not fertile or even until her death is common in equine industry but the fertility decreases as the mare age increases. Embryo loss due to reduced embryo quality is partly accountable for this observation. Here, the effect of mare’s age on blastocysts’ gene expression was explored. Day 8 post-ovulation embryos were collected from multiparous young (YM, 6-year-old, N = 5) and older (OM, > 10-year-old, N = 6) non-nursing Saddlebred mares, inseminated with the semen of one stallion. Pure or inner cell mass (ICM) enriched trophoblast, obtained by embryo bisection, were RNA sequenced. Deconvolution algorithm was used to discriminate gene expression in the ICM from that in the trophoblast. Differential expression was analyzed with embryo sex and diameter as cofactors. Functional annotation and classification of differentially expressed genes and gene set enrichment analysis were also performed.

**Results:** Maternal aging did not affect embryo recovery rate, embryo diameter nor total RNA quantity. In both compartments, the expression of genes involved in mitochondria and protein metabolism were disturbed by maternal age, although more genes were affected in the ICM. Mitosis, signaling and adhesion pathways and embryo development were decreased in the ICM of embryos from old mares. In trophoblast, ion movement pathways were affected.

**Conclusions:** This is the first study showing that maternal age affects gene expression in the equine blastocyst, demonstrating significant effects as early as 10 years of age. These perturbations may affect further embryo development and contribute to decreased fertility due to aging.

## Introduction

Unlike other farm animals, the main purpose for the use of horses is athletic performance and leisure. In many cases, mares produce foals regularly to remain profitable after their sport career so that their reproductive career may last until advanced age, even though mares’ fertility declines with age (for review ^1^). Indeed, in Thoroughbreds, pregnancy rates per estrous at Day 42 post-ovulation were shown to be reduced from 63.0% in young mares to 58.5% in mares older than 9 years of age, and to reach 43.8% in mares older than 18 years old ^2^. It is difficult to determine an age limit defining when a mare is considered too old for breeding as aging is a progressive process during the life. One study on 535,746 fertility records in mixed breeds in France, however, suggested that if a 10% of fertility decline is chosen as cutoff value for retiring breeding mares, then mares older than 8 should not be bred ^3^.

Embryo loss seems to be the principal cause for the fertility decline in old mares. Using pregnancy data obtained from 898 cycles in Thoroughbred mares over period of 11 years, the cutoff for embryo mortality to reach values above average was calculated to be 10 years of age ^4^. In practice, embryo loss frequency is generally observed between the first (14 - 22 days) and the second (42 - 45 days) pregnancy check in old than in young mares ^2,4,5^. Embryo loss may, however, be unobserved in earlier stages. Indeed, although experimental oviductal flushes on Day 2 post-ovulation showed that the fertilization rate was similar between young and fertile mares *vs* old and subfertile mares (respectively, 10/14 *vs* 11/14 embryos recovered per group on Day 2), when the same mares were inseminated during another estrus period, the pregnancy rate was significantly higher at 14 days post ovulation in young and fertile mares compared to older ones (respectively, 12/15 *vs* 3/15 pregnant mares on Day 14 by ultrasonography) ^6^.

Although uterine degeneration ^7–9^ is observed in old mares, oocyte quality seems to play a major role in the observed age-related fertility decrease: the transfer of Day 4 embryos collected from old and subfertile mares resulted in lower pregnancy rates at 14 days post-ovulation than when embryos collected from young and fertile mares were transferred. This remained true when only the subset of good quality embryos was considered ^10^.

Controversial effects of maternal age on oocyte morphology have been observed in several studies ^11–15^ but there is a clear effect of maternal age on oocyte quality and function. Old mares’ oocytes are less likely to reach metaphase II after *in vitro* maturation than oocytes from younger mares ^11^. Alterations of the expression of several genes have been also observed during *in vivo* and *in vitro* oocyte maturation ^16,17^. Moreover, after *in vitro* maturation, higher spindle instability is observed in old versus young mares’ oocytes, resulting in increased risks of chromosome misalignment leading to embryo aneuploidy ^18–20^. Controversial effects of mare age on mitochondria degeneration have been also reported ^16,21–23^.

The effect of maternal age on embryo size is not so clear, as studies demonstrated a reduction of embryo diameter in old mares at several developmental stages ^24–28^ while others did not detect any effect ^9,13,25,29^ of maternal aging on embryo size. In these studies, however, the developmental stage was not uniform and the method to measure embryo size could be different (direct measurement or using ultrasonography). Furthermore, the size of the equine embryo is highly variable for the same gestational age ^30^. The only study that directly measured more than 200 Day 7 to 10 embryos as part of an embryo transfer program observed that maternal aging induced a reduction in embryo size (in average, more than 740µm in diameter for embryos produced by mares younger than 15 *vs* less than 620µm in diameter for embryos produced by mares older than 16) ^31^.

Maternal age, however, seems to affect embryo morphology from the earliest stages of development. Embryos collected *in vivo* on Day 1.5 post ovulation, indeed, had the same number of cells, but a poorer morphology grade (uniformity of cell size and texture, imperfections and abnormalities) when they had been collected from mares older than 15 years of age compared to young ones ^32^. In addition, at Day 3 post ovulation, embryos collected from mares older than 20 years of age combined poorer morphology grade and reduced cell number ^32^. In contrast, in other studies, no difference according to maternal age was observed in terms of number, size or texture of blastomeres of embryos collected in oviducts on Days 2 and 4 post ovulation ^6,10^. Finally, at blastocyst stage, a higher incidence of morphological abnormalities was observed in embryos that were collected from subfertile and/or 18 years old mares compared to 4 years old nulliparous mares ^24^. Only a few studies have examined the effect of maternal age on equine embryo quality. As observed in oocytes, metabolic function and capacity were reduced in Day 2 embryos collected from old mares (≥20 years old) compared to those collected from young mares (≤14 years old) ^22^. Moreover, a reduced quantity of mitochondrial DNA, but not genomic DNA, has been reported in Day 7 blastocysts from mares older than 16 years of age compared to younger counterparts, suggesting a reduction in mitochondria/cell ratio in old mares’ blastocysts ^23^. Taken together, these alterations could lead to the alterations in embryo mobility and/or day of fixation that have been observed in some studies ^9,28^.

Altogether, these data indicate that the quality of embryos is reduced in older mares. Considering the Developmental Origins of Health and Diseases (DOHaD) ^33^, dam age could thus also affect feto-placental and post-natal growth as well as long-term offspring health. To the authors’ knowledge, the effect of maternal age on gene expression in the equine embryo, however, has not yet been explored.

The aim of this study was to determine the effect of maternal age in the mare on embryo gene expression at the blastocyst stage. Young (<10 years) and older (>10 years) multiparous mares were inseminated with the same stallion. Blastocysts were collected on Day 8 post-ovulation and bisected to separate the pure trophoblast (TE_part) from the inner cell mass enriched hemi-embryo (ICMandTE). Gene expression was analyzed by RNA-seq in each compartment. Deconvolution was used to isolate gene expression of inner cell mass from trophoblast cells in ICMandTE part.

## Methods

### Ethics

The experiment was performed at the experimental farm of IFCE (research agreement C1903602 valid until March 22, 2023). The protocol was approved by the local animal care and use committee (“Comité des Utilisateurs de la Station Expérimentale de Chamberet”) and by the regional ethical committee (“Comité Régional d’Ethique pour l’Expérimentation Animale du Limousin”, approved under N° C2EA - 33 in the National Registry of French Ethical Committees for animal experimentation) under protocol number APAFIS#14963-2018050316037888 v2. All experiments were performed in accordance with the European Union Directive 2010/63EU. The authors complied with the ARRIVE guidelines.

### Embryo collection

Thirty-one non-nursing multiparous Saddlebred mares (French Anglo-Arabian and Selle Français breeds) were included in this study. Multiparous mares were defined as dams that had already foaled at least once. Mares were bred and raised in the “Institut Français du Cheval et de l’Equitation” experimental farm (Chamberet, France, 45° 34’55.17”N, 1°43’16.29”E, 442m). During the experimental protocol, mares were managed in herds in natural pastures 24h/day with free access to water. The experiments took place from April 1^st^ to May 23^rd^, 2019. All mares remained healthy during this period.

Mares were allocated to 2 groups according to their age: young mares (YM, < 10 years, n = 11) and older mares (OM, ≥ 10 years, n = 20). The age threshold was defined in accordance with the literature ^3,4^. Before experimentation, mare’s withers’ height was measured. During the experimentation, mares were weighted and mares’ body condition score (BCS, scale 1 to 5) was determined by one trained operator ^88^.

The mares’ estrous period was monitored routinely by ultrasound with a 5MHz trans-rectal transducer. During estrus, ovulation was induced with a single injection of human chorionic gonadotropin (i.v.; 750 - 1500IU; Chorulon® 5000; MSD Santé animale, France) as soon as one ovarian follicle > 35mm in diameter was observed, together with marked uterine edema. Ovulation usually takes place within 48h, with > 80% occurring 25 to 48h after injection ^89,90^. At the same time, mares were inseminated once with fresh or fresh overnight cooled semen containing at least 1 billion motile spermatozoa from a single fertile stallion. Ovulation was confirmed within the next 48 hours by ultrasonography.

Embryos were collected by non-surgical uterine lavage using prewarmed (37°C) lactated Ringer’s solution (B.Braun, France) and EZ-Way Filter (IMV Technologies, France) 10 days after insemination, i.e. approximately 8 days post ovulation. Just after embryo collection, mares were treated with luprotiol an analogue of prostaglandin F2α (i.m; 7.5 mg; Prosolvin, Virbac, France).

The aim of the embryo collection was to obtain 5 embryos/group with each embryo coming from a different mare. Therefore, some mares that failed to produce an embryo at their first attempt had been bred again.

### Embryo bisection and RNA extraction

Collected embryos were immediately observed using a binocular magnifier and photographed with a size standard. Embryo diameter was subsequently determined on photographs using ImageJ® software (version 1.52a; National Institutes of Health, Bethesda, MD, USA). Embryos were washed 4 times in commercially available Embryo holding medium (IMV Technologies, France) at 34°C and then bisected with a microscalpel under binocular magnifying glass to separate trophoblast part, at this stage composed of trophectoderm and endoderm (TE_part) from inner cell mass (composed of epiblast cells) enriched hemi-embryos (ICMandTE) (Figure 1). Each hemi-embryo was transferred to an Eppendorf DNA LoBind tube (Eppendorf, Montesson, France) with extraction buffer (PicoPure RNA isolation kit, Applied Biosystems, France) and incubated for 30 min at 42°C prior to storage at - 80°C.

**Figure 1:**
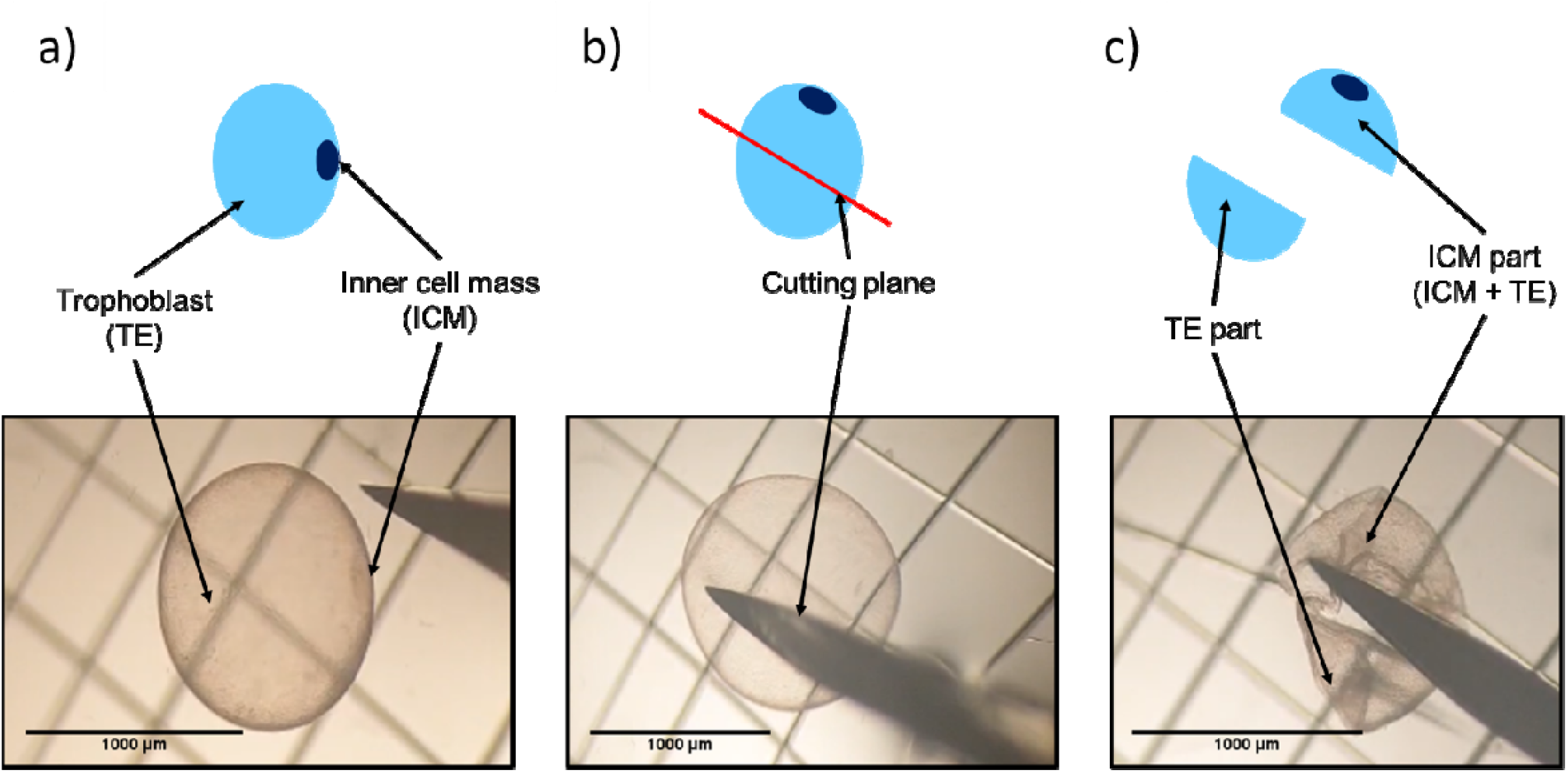
Bisection of equine embryos at 8 days post-ovulation into an ICMandTE and a TE_part. a) Step 1: Identification of the ICM and the TE; b) Step 2: Definition of the cutting plane to isolate the ICM in one of the parts; c) Step 3: Cutting of the embryo and separation of the two parts. ICMandTE: inner cell mass + trophoblast; TE part: pure trophoblast. NB: Here the trophoblast represents trophectoderm + endoderm and ICM is composed of epiblast cells.

RNA was extracted from each hemi-embryo following the manufacturer’s instructions using PicoPure RNA isolation kit (PicoPure RNA isolation kit, Applied Biosystems, France), which included a DNAse treatment. RNA quality and quantity were assessed with the 2100 Bioanalyzer system using RNA 6000 Pico or nano kit (Agilent Technologies, France) according to the manufacturer’s instructions (Supplementary Figure S1).

### RNA sequencing

Five nanograms of total RNA were mixed (Supplementary Figure S1) with ERCC spike-in mix (Thermofisher Scientific, France) according to manufacturer recommendations. Messenger RNAs were reverse transcribed and amplified using the SMART-Seq V4 ultra low input RNA kit (Clontech, France) according to the manufacturer recommendations. Nine PCR cycles were performed for each hemi-embryo. cDNA quality was assessed on an Agilent Bioanalyzer 2100, using an Agilent High Sensitivity DNA Kit (Agilent Technologies, France). Libraries were prepared from 0.15 ng cDNA using the Nextera XT Illumina library preparation kit (Illumina, France). Libraries were pooled in equimolar proportions and sequenced (Paired end 50-34 pb) on NextSeq500 instrument, using a NextSeq 500 High Output 75 cycles kit (Illumina, France) (Supplementary Figure S1). Demultiplexing was performed with bcl2fastq2 version 2.2.18.12 (Illumina, France) and adapters were trimmed with Cutadapt version 1.15 ^91^. Only reads longer than 10pb were kept.

### Embryo sexing

Embryo sex was determined for each embryo in order to take in consideration embryo sex in the statistical analysis as sexual dimorphism was observed in blastocysts in other species ^92^. The expression of *X Inactive Specific Transcript* (*XIST*) was analyzed. *XIST* is a long noncoding RNA from chromosome X responsible for X inactivation when two X chromosomes are present, meaning that only female embryos express *XIST. XIST* was shown to be expressed in equine female blastocysts on Day 8 post ovulation ^35,38^. *XIST* is not annotated in the available equine Ensembl 3.0.99 or NCBI annotation. Thus, a *de novo* transcript assembly was made using StringTie version 2.1.2 ^93^ following manufacturer instructions to identify *XIST* in equine embryo samples (Supplementary Figure S1). The genome annotation file output was aligned on Ensembl 99 EquCab3.0 assembly using IGV version 2.7.2 ^94^. As *XIST* position and length on X chromosome has been previously determined ^35^ and as *XIST* is annotated in humans, the most similar contig was selected. The obtained *XIST* sequence was 22,600 bp long and composed of 8 exons and 7 introns. *XIST* sequence was added to FASTA files from Ensembl 99 EquCab3.0 assembly and to the genome annotation file from Ensembl 99 EquCab3.0 assembly as a unique fictitious chromosome with one gene composed of one transcript.

*XIST* expression differed between individuals but was relatively close in quantification between related hemi-embryos. Therefore, seven embryos were determined as females (4 in the YM group and 3 in the OM group) while 4 were considered as males (1 in the YM group, and 3 in the OM group).

### RNA mapping and counting

Alignment was performed using STAR version 2.6 ^95^ (2-pass mapping with the genome annotation file using the following parameters: --alignSoftClipAtReferenceEnds No --alignEndsType EndToEnd -- alignIntronMax 86545 --alignMatesGapMax 86545) on previously modified Ensembl 99 EquCab3.0 assembly and annotation. Genes were then counted with featureCounts ^96^ from Subreads package version 1.6.1 (parameters: -p -B -C -M) (Supplementary Figure S1).

### Data analysis

All statistical analyses were performed by comparing OM to YM (set as reference group) using R version 4.0.2 ^97^ on Rstudio software version 1.3.1056 ^98^.

### Embryo recovery rate, embryo diameter and total RNA content analysis

Embryo recovery rate (ERR) per mare was calculated as number of attempts with at least one embryo collected/total number of attempts. ERR per ovulation was determined as number of collected embryos/total number of observed ovulations. They were analyzed using the Exact Fisher test to determine if age influences ERR.

For total RNA content analysis, as embryos were bisected without strict equality for each hemi-embryo, a separate analysis of ICMandTE and TE_part RNA quantities would not be meaningful. Thus, ICMandTE and TE_part RNA quantities were initially summed up and analyzed using a mixed linear model from nlme package version 3.1-148 ^99^, followed by 1000 permutations using PermTest function from pgirmess package version 1.6.9 ^100^. Maternal age and embryo sex were considered as random effects.

Embryo diameter was analyzed with a fixed effects linear model of nlme package version 3.1-148 ^99^ including maternal age and embryo sex, followed by 1000 permutations using PermTest function from pgirmess package version 1.6.9 ^100^. Variables were kept in models when statistically significant differences were observed. Differences were considered as significant for p < 0.05.

Moreover, the relation between embryo size and total RNA quantity/embryo was studied and represented using the nonlinear regression analysis (fitness to exponential growth equation) in GraphPad Prism software 8.0.1 for Windows (Graphpad Software, San Diego, California USA, www.graphpad.com).

### Deconvolution of gene expression in ICMandTE using DeMixT

A filter on counts obtained with featureCounts was applied: all genes with more than 3 non null count values in at least one group (OM or YM) per hemi-embryo (ICMandTE or TE_part) were conserved. The ICMandTE hemi-embryo was composed of both trophoblast and inner cell mass in unknown relative proportions. In order to estimate the relative gene expression of both cell types, the DeMixT R package version 1.4.0 was used ^101,102^. Starting from a dataset obtained from one cell type, DeMixT estimates the proportion of cells in the mixed samples and performs a deconvolution algorithm on gene expression. TE_part (reference cells) and ICMandTE (mixed cells type) datasets were thus analyzed using the following options: niter=10; nbin=50; if.filter=TRUE; ngene.selected.for.pi=1500; ngene. Profile.selected=1500; nspikein=0. Since the deconvolution algorithm cannot be performed in the presence of null values, the value “1” was added to all count values in ICMandTE and TE_part data tables. Moreover, variance in reference cell dataset (TE_part) must not be null and gene with null variance had been removed prior to the deconvolution. Output datasets were DeMixT_ICM_cells and DeMixT_TE_cells, corresponding to the deconvoluted gene expression in ICM cells and TE cells of ICMandTE, respectively.

### Quality checks for deconvolution

At the end of deconvolution, a quality check is automatically performed by the analysis: for every gene, if the difference of average of deconvoluted expression of reference cells in mixed samples (here, gene expression of TE cells in ICMandTE dataset, *i*.*e*., DeMixT_TE_cells) and average of expression of reference cells (here, gene expression of TE cells in TE_part) is > 4, then, the deconvolution for concerned genes is not reliable. Therefore, genes in this case are filtered out.

Moreover, outputs of DeMixT_ICM_cells *vs* DeMixT_TE_cells, DeMixT_ICM_cells *vs* TE_part and ICMandTE *vs* TE_part were compared with Deseq2 version 1.28.1 ^103^ to confirm that the deconvolution was effective at separating gene expression. Moreover, several genes proper to ICM and TE cells in equine embryos were identified using literature search ^35^. Their raw and deconvoluted expressions were analyzed to check the reliability of deconvolution. Results of these analyses were represented through manually drawn Venn diagrams as well as principal component analysis (PCA) graphics of individuals, using FactoMineR version 2.4 ^104^, ggplot2 version 3.3.3 ^105^ and factoextra version 1.0.7 ^106^.

### Maternal age comparison for gene expression

For the comparison of maternal age, ICMandTE, DeMixT_ICM_cells and TE_part datasets were used and a filter was applied on datasets: all genes with an average expression equal or above 10 counts in at least one group (OM or YM) per hemi-embryo (ICM or TE) were conserved. Differential analyses were performed with Deseq2 version 1.28.1 ^103^ with YM group as reference, without independent filtering. Genes were considered differentially expressed (DEG) for FDR < 0.05 after Benjamini-Hochberg correction (also known as false discovery rate, FDR). As ovulation was checked only every 48h and because embryos growth is exponential in the uterus, embryo diameter was considered as a cofactor in the model as well as embryo sex. Moreover, PCAs using factominer version 2.4 ^104^on R software followed with a Partial redundancy analysis using rda function from vegan package version 2.5-7 ^107^ were performed. For these analyses, embryo size was considered as a factor of two modalities: embryos < 1100µm were considered as small and > 1100µm were considered as large. Results were represented with a biplot graphic using the package factoextra version 1.0.7 ^106^. The combined analyses were used to represent and determine the specific contribution of each of the two factors in the total variability observed in DEGs. The objective was to verify that the age of the mare was the main factor explaining the difference between the DEGs and not the differences in embryo size. On biplot graphics, only the 20 variables, *i*.*e*., genes, that contributed the most in axis construction, were represented.

Equine Ensembl IDs were converted into Human Ensembl IDs and Entrez Gene names using gorth function in gprofiler2 package version 0.1.9 ^108^. Genes without Entrez Gene names using gprofiler2 were manually converted when Entrez Gene names were available, using Ensembl web search function ^109^.

Moreover, DEG were analyzed for classification, over- and under-representation (FDR < 0.05) using Fisher’s exact test corrected with FDR and using the database Gene Ontology (GO) Biological Process complete on PANTHER version 16.0 web-based software ^110^. Only the most specific subclass pathways, provided by PANTHER’s automated hierarchy sorting, were conserved and represented.

### Gene set enrichment analyses (GSEA)

The count values were log transformed using RLOG function of DESeq2 version 1.28.1. Gene set enrichment analyses (GSEA) were performed on expressed genes using GSEA software version 4.0.3 (Broad Institute, Inc., Massachusetts Institute of Technology, and Regents of the University of California) ^111,112^ to identify biological gene sets disturbed by age. GSEA was performed using the following parameters: 1000 gene set permutations, weighted enrichment analysis, gene set size between 15 and 500, signal-to-noise metrics. Molecular Signatures Database ^113^ version 7.1 (C2: KEGG: Kyoto Encyclopedia of Genes and Genomes, C5: BP: GO biological process) were used to identify most perturbed pathways. Pathways were considered significantly enriched for FDR < 0.05. When normalized enrichment score (NES) was positive, the gene set was enriched in the OM group while when NES was negative, the gene set was enriched in the YM group.

Enriched terms from GO BP and KEGG database were represented using SUMER analysis from SUMER R package version 1.1.5 and using FDR q-values ^114^. First, SUMER analysis reduces redundancy using weighted set cover algorithm and then clusters the results using affinity propagation algorithm. The weighted set cover algorithm performs a selection of the fewest number of gene sets that include all genes associated with the enriched sets, giving priority for the most significant terms. Then, the affinity propagation algorithm groups similar gene sets and identifies one representative gene set. This algorithm will repeat this clustering to obtain the most representative gene sets. Results were represented with graph modified using Cytoscape version 3.8.2 ^115^. Gene sets were represented by nodes and the gene set size was represented by the size of the node. Node shape represented the gene set database (GO BP or KEGG). Blue nodes represented gene sets enriched in YM (NES < 0) while red nodes represented gene sets enriched in OM (NES > 0). Edge width represents the level of connection between representative gene sets (thinner edges represent the first clustering while thicker edges represent the second clustering of the affinity propagation algorithm)

## Data availability statement

The datasets generated during and/or analyzed during the current study are available in the GEO repository,

[accession: GSE162893; https://www.ncbi.nlm.nih.gov/geo/query/acc.cgi?acc=GSE162893].

## Results

Altogether, 36 uterine flushings were performed (12 in YM <10 years old and 24 in OM >10 years old) to obtain one embryo per mare. For 5 mares, after a negative embryo collection, a second attempt was performed. Characteristics of mares are detailed in Table 1. One YM and 3 OM had a double ovulation but no twin embryo was obtained. Embryo recovery rates (ERR) per mare and per ovulation were, respectively, 42% and 39% in YM and 25% and 22% in OM, with no significant difference between groups (p = 0.45 for ERR/mare and p = 0.46 for ERR/ovulation). All embryos were expanded blastocysts, grade I or II according to the embryo classification of McKinnon and Squires ^34^. Embryo diameter ranged from 409 µm to 2643 µm, with no effect of group on embryo diameter (p = 0.97). RNA yield per embryo ranged from 16 ng to 2915.5 ng and was not related to mares’ age (p = 0.92). There is a clear exponential relation between RNA yield/embryo and embryo size (Figure 2a). The median RNA Integrity Number (RIN) was 9.8 (8.7 - 10 range). Between 29.4 and 69.5 million reads per sample were obtained after trimming. On average, 83.61% of reads were mapped on the modified EquCab 3.0 using STAR and 66.66% of reads were assigned to genes by featureCounts.

**Table 1:** Mares’ characteristics at embryo collection time.

| Characteristics | Young mares (YM)<br>(n = 11) |  | Old mares (OM)<br>(n=20) |  |
| --- | --- | --- | --- | --- |
|  | Total | With embryo (n = 5) | Total | With embryo (n = 6) |
| Breed | AA n = 9; SF n =1; SB n=1 | AA | AA n = 13; SF n = 7 | AA n = 4; SF n = 2 |
| Age (*in years*) | 6.00 ± 0.00 | 6.00 ± 0.00 | 11.95 ± 2.11 | 12.00 ± 2.90 |
| Parity (*number of foalings*) | 1.00 ± 0.00 | 1.00 ± 0.00 | 3.20 ± 1.15 | 3.17 ± 1.47 |
| Weight (*in kg*) | 536.62 ± 44.12 | 524.02 ± 61.39 | 589.35 ± 57.34 | 598.19 ± 76.83 |
| BCS (*scale 1-5*) | 2.16 ± 0.28 | 2.3 ± 0.33 | 2.61 ± 0.32 | 2.5 ± 0.59 |
| Withers' height (*in cm*) | 157.95 ± 3.74 | 157.40 ± 5.41 | 161.21 ± 4.23 | 162.00 ± 6.16 |

All mares were multiparous.
AA: Anglo Arab or Anglo-Arabian type; SF: Selle Français; SB: Saddlebred; BCS: Body Condition Score.
Age, parity, weight, BCS and height are presented as mean ±SD

**Figure 2:**
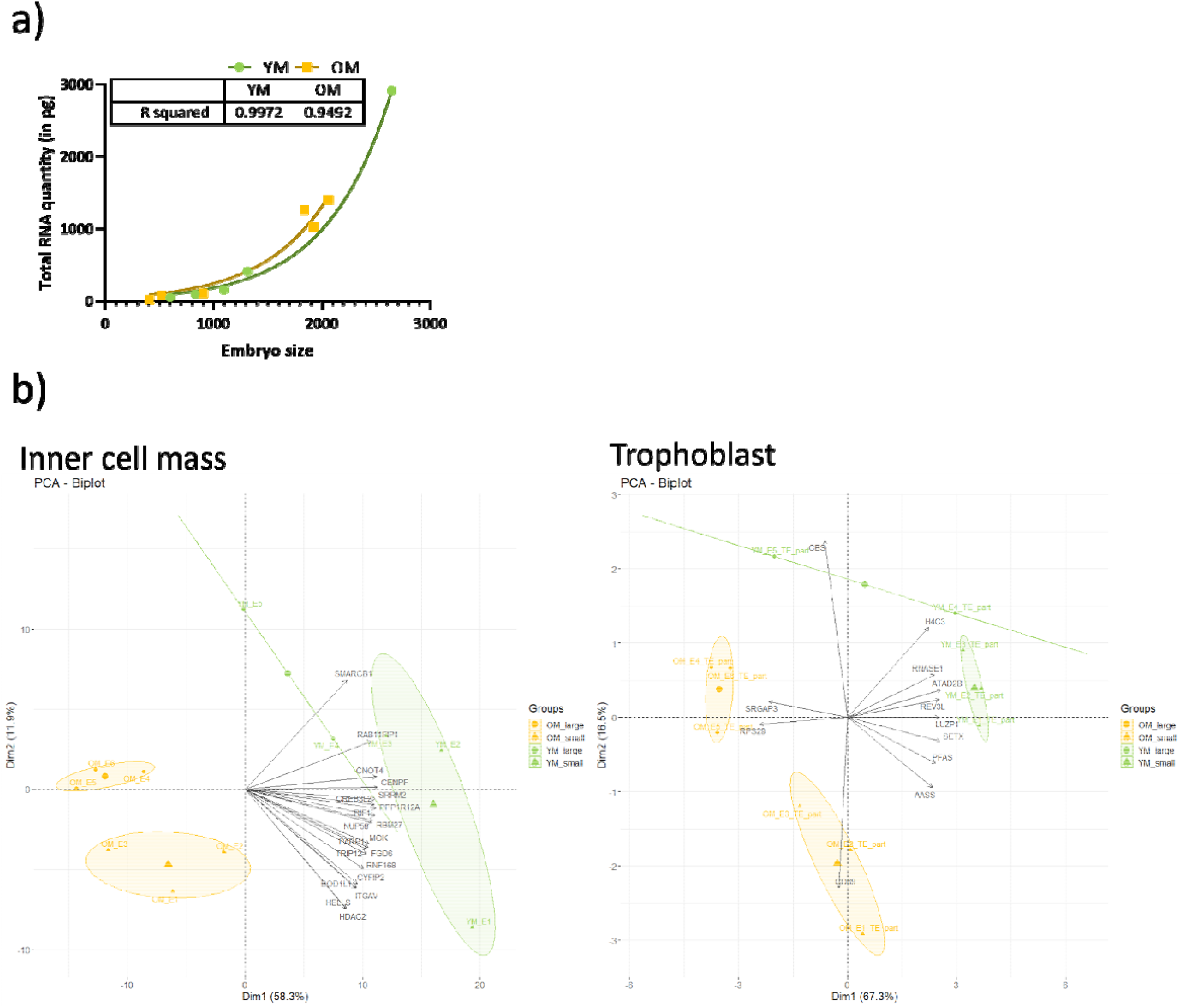
Effect of embryo size on RNA yield and gene expression in equine embryos a) Green circles represent embryos from young mares (YM) and yellow squares represent embryos from old mares (OM). The used relation is exponential. b) Biplot graphics representing principal component analyses of differentially expressed genes in inner cell mass and trophoblast of embryos collected in YM or OM. Embryo were grouped according to maternal age (YM or OM) and their diameter (< 1100µm = small; > 1100µm = large). The top 20 variables contributing to the axis formation were represented.

After keeping genes with less than 3 non null count values in at least one group (OM or YM) per hemi-embryo (ICMandTE or TE_part), 16,741 genes were conserved for deconvolution. In addition, twenty-seven genes were removed because their variance was null in the TE_part. However, for these genes, the mean counting in ICMandTE samples was above 10 counts, excepted for one which had a mean expression of 34 counts. On the other hand, this gene was only expressed in 5 ICMandTE over the 10 counts threshold.

During the analysis, 3 additional genes were excluded because the deconvolution quality for these genes was not sufficient. Therefore, at the end of deconvolution algorithm, 16,711 genes were available for differential analysis.

Before deconvolution, 341 genes were differentially expressed (FDR < 0.05) between the ICMandTE and the TE_part (Figure 3a). After deconvolution, comparison between DeMixT_ICM_cells and DeMixT_TE_cells yielded 7,565 differentially expressed genes while the comparison DeMixT_ICM_cells *vs* TE_part yielded 6,085 differentially expressed genes, with 5,925 in common (72%). Moreover, except one of the initially 341 differentially expressed genes before deconvolution, were also identified as differentially expressed in both post-deconvolution analyses. On the PCA graphic of individuals, ICMandTE and TE_part were not really separated before deconvolution (Figure 3b). DeMixT_TE_cells and TE_part were partly superposed, suggesting that datasets before and after deconvolution have a similar global gene expression; whereas the DeMixT_ICM_cells group is clearly separated from both on Axis 2 (14.3% of variance), indicating that the deconvolution effectively enabled the separation of gene expression in the two cell types.

**Figure 3:**
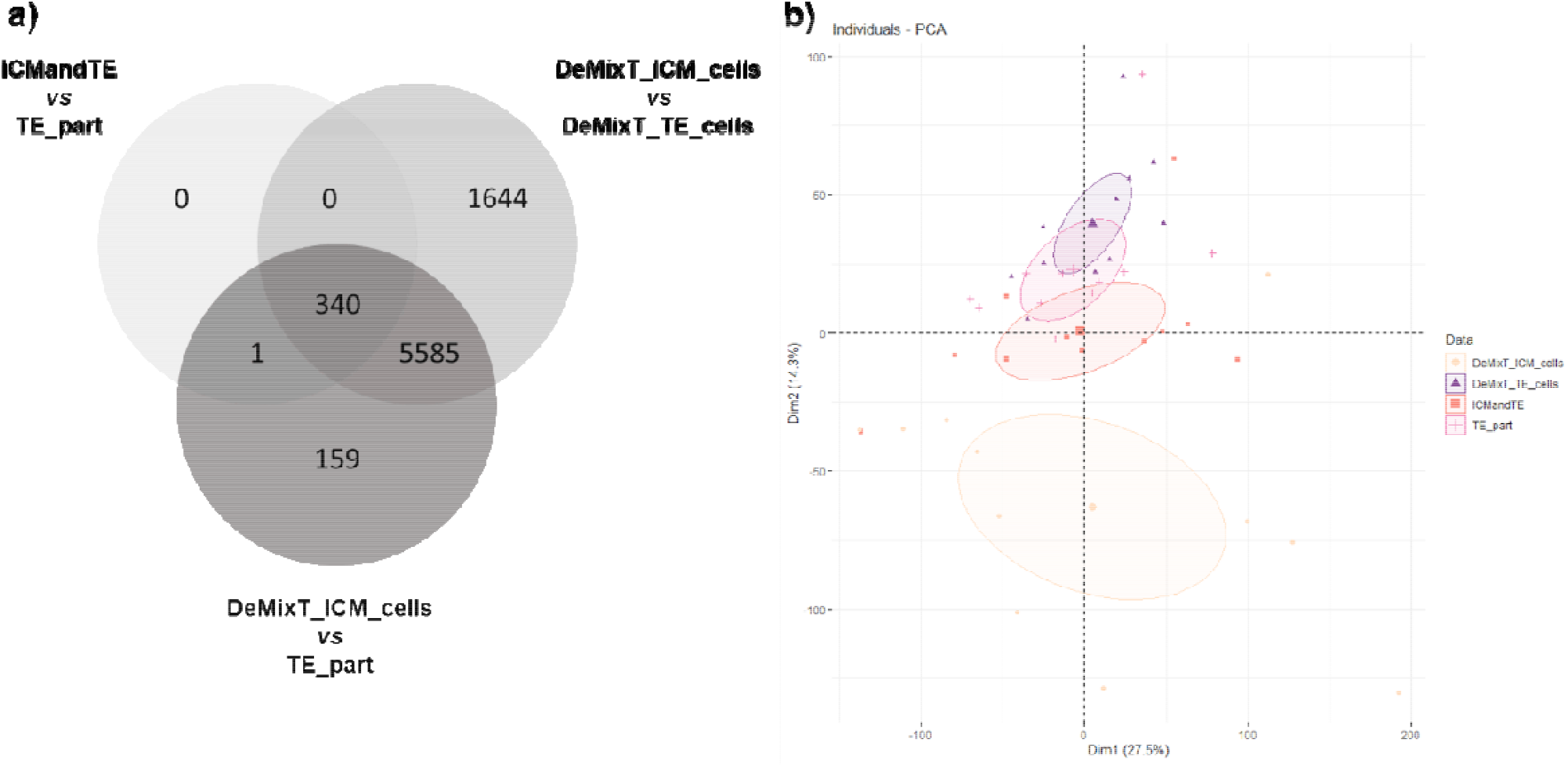
Gene expression in ICM and TE before and after deconvolution using DeMixT A) Venn diagrams of the differential gene expression in ICMandTE *vs* TE part (before deconvolution), DeMixT_ICM_cells *vs* DeMixT_TE_cells (after deconvolution) and DeMixT_ICM_cells *vs* TE_part (gene expression of ICM after deconvolution *vs* gene expression in TE_part without deconvolution); B) Principal Component Analysis of gene expression of DeMixT_ICM_cells, DeMixT_TE_cells, ICMandTE and TE part datasets. Deconvolution was used to isolate gene expression of ICM and TE cells in ICMandTE hemi-embryos. ICMandTE: inner cell mass + trophoblast; TE part: pure trophoblast. Here trophoblast represents trophectoderm + endoderm.

Six of the 12 genes previously identified as more expressed in ICM ^35^ were also more expressed in ICMandTE *vs* TE_part comparison (Table 2). After deconvolution (comparison DeMixT_ICM_cells *vs* TE_part), 3 more genes previously identified were also differentially expressed. However, for two of them (POU Class 5 Homeobox 1, *POU5F1* and Undifferentiated Embryonic Cell Transcription Factor 1, *UTF1*), their expression was increased in TE_part after deconvolution. In TE, any genes previously identified were differentially expressed in the comparison ICMandTE *vs* TE_part, *i*.*e*., before deconvolution. After deconvolution, the expression of 2 genes was increased in TE_part compared to DeMixT_ICM_cells.

**Table 2:** Comparison of selected genes expression before and after deconvolution Gene expressions were obtained from RNA of 11 equine embryos bissected in two hemi-embryos: one part is composed only of trophoblast (TE), TE_part, while the other part is composed of TE and inner cell mass (ICM), ICMandTE. As it is impossible to estimate the proportion of each cells in ICMandTE, deconvolution algorithm (package DeMixT) was used to estimate gene expression of these different kind of cells. DeMixT_ICM_cells dataset is the deconvoluted gene expression of ICM cells from ICMandTE. Log2 fold change (log2FC) and padj (adjusted p-value with Benjamini-Hochberg correction) were obtained with Deseq2 package. TE_part is the reference group in both analyses: when log2 fold changes (log2FC) > 0, gene is more expressed in ICMandTE or DeMixT_ICM_cells while when log2FC < 0, gene is more expressed in TE_part. Green is used to represent gene differentially expressed in the present study. Orange is used to represent gens that have been previously identified as predominant in the ICM ^35^ but which are identified here as predominant in the TE.

|  | Gene name | Ensembl ID | ICMandTE vs TE_part |  | DeMixT_ICM_cells vs TE_part |  |
| --- | --- | --- | --- | --- | --- | --- |
|  |  |  | log2FC from DeSeq2 | padj | log2FC from DeSeq2 | Padj |
| **ICM** | **SOX2** | ENSECAG00000010653 | 4.49 | 0.02 | 6.49 | 1.7E-06 |
|  | **NANOG** | ENSECAG00000012614 | 4.24 | 1.0E-07 | 5.98 | 2.4E-15 |
|  | **SPP1** | ENSECAG00000017191 | 4.02 | 8.4E-09 | 5.74 | 5.1E-16 |
|  | **LIN28B** | ENSECAG00000020994 | 3.23 | 1.3E-10 | 5.00 | 4.7E-22 |
|  | **SMARCA2** | ENSECAG00000024187 | 0.62 | 0.97 | 1.25 | 0.07 |
|  | **POU5F1 (OCT4)** | ENSECAG00000008967 | 0.60 | 0.009 | -2.04 | 4.6E-07 |
|  | **ID2** | ENSECAG00000008738 | 0.44 | 0.29 | 1.24 | 9.0E-05 |
|  | **DNMT3B** | ENSECAG00000012102 | 0.35 | 0.24 | 0.8 | 0.06 |
|  | **DPPA4** | ENSECAG00000013271 | 0.35 | 0.04 | 1.14 | 1.8E-05 |
|  | **SALL4** | ENSECAG00000018533 | 0.10 | 1 | 0.60 | 0.01 |
|  | **KLF4** | ENSECAG00000010613 | -0.04 | 1 | -0.81 | 0.39 |
|  | **UTF1** | ENSECAG00000039888 | -0.13 | 1 | -2.21 | 7.3E-05 |
| **TE** | **TFAP2A** | ENSECAG00000017468 | -0.17 | 0.46 | -0.30 | 0.004 |
|  | **CDX2** | ENSECAG00000027754 | -0.14 | 0.77 | -0.23 | 0.07 |
|  | **ELF3** | ENSECAG00000014608 | -0.13 | 1 | -0.19 | 0.27 |
|  | **GATA2** | ENSECAG00000016768 | -0.13 | 1 | -0.29 | 0.048 |
|  | **GATA3** | ENSECAG00000024574 | -0.10 | 0.87 | -0.05 | 0.64 |
|  | **TEAD4** | ENSECAG00000011303 | -0.10 | 0.96 | 0.00 | 1 |
|  | **FREM2** | ENSECAG00000020410 | 0.07 | 1 | 0.19 | 0.44 |

For further analysis, DeMixT_ICM_cells and TE_part gene expression datasets are presented. Nevertheless, the comparison of maternal age on ICMandTE dataset (graphics, differential gene expression analysis and GSEA) was also performed and this analysis is available in Supplementary Figure S2 and supplementary Tables S1 and S2.

After filtering out of genes with an average expression < 10 counts/maternal age group/hemi-embryo, 13,735 genes were considered as expressed in the ICM cells from OM or YM embryos including 255 differentially expressed genes (227 downregulated and 28 upregulated in OM) (Figure 4a and Supplementary Table S3). Only 218 and 12 genes out of the down- and upregulated genes, respectively, were associated to a known protein in human. These downregulated genes in ICM of embryos from OM mainly belonged to cellular process, metabolic process and biological regulation in GO biological process gene sets (Figure 4a). From downregulated genes in OM ICM, 84 pathways were statistically overrepresented in GO Biological process leading to 16 most specific subclass pathways according to hierarchy sorting by PANTHER (Figure 4b). Overrepresented pathways in downregulated genes in OM ICM were linked to cell division, embryo morphogenesis, histone methylation, protein modification processes and cytoskeleton organization.

**Figure 4:**
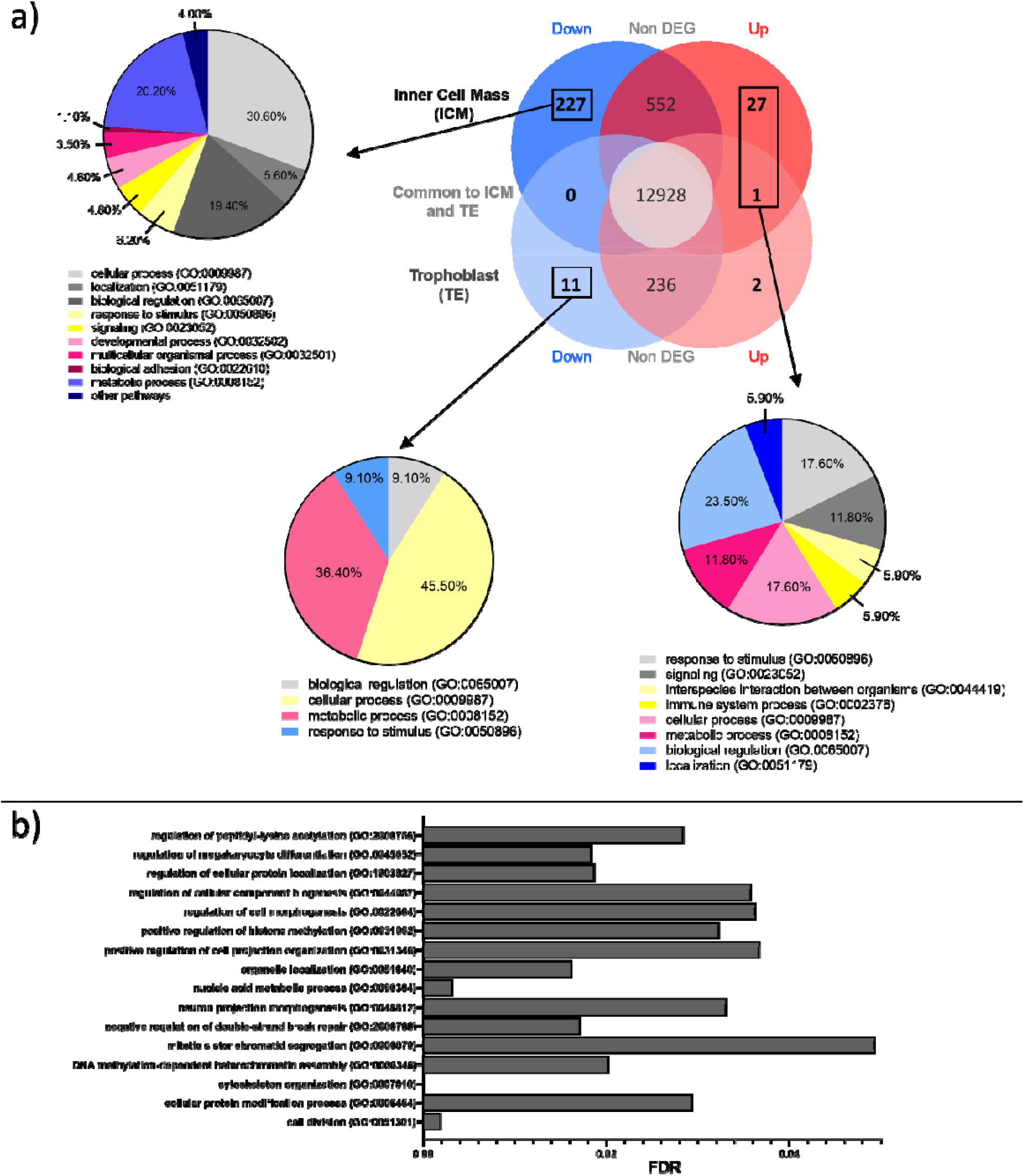
Analysis of differentially expressed genes (DEG) in embryos according to maternal age A) representation of down (blue) and upregulated (red) DEG in ICM (from DeMixT_ICM_cells data obtained after deconvolution of ICMandTE using DeMixT R package ^101,102^) and TE (from TE_part dataset) of embryos from OM *vs* YM. Functional classifications (for down regulated genes in ICM and TE and for up-regulated genes in ICM) using GO Biological Process obtained with PANTHER online software are presented as pie charts. For upregulated DEG in TE, no functional classification was made. B) Bar chart presenting p-adjusted of most specific subclass pathways provided by PANTHER online software after statistical overrepresentation test (Fisher’s Exact test and Benjamini-Hochberg correction) with the Human GO Biological Process annotation on down-regulated DEGs in ICM. DEG: Differentially Expressed Genes (FDR < 0.05); TE: Trophoblast; ICM: Inner Cell Mass; OM: Old multiparous mares; YM: young multiparous mares

Upregulated differentially expressed genes (DEG) in OM ICM were mainly part of biological regulation, response to stimulus, cellular process, signaling and metabolic process in GO biological process gene sets (Figure 4a). No pathway was statistically overrepresented in upregulated genes in ICM of embryos from old mares using PANTHER overrepresentation testing.

In the TE, 13,178 genes were considered as expressed in OM or YM. Fourteen were differentially expressed (Supplementary Table S4) with 11 genes being downregulated and 3 being up regulated in OM (Figure 4a). Only 12 genes were associated to a known protein in human. Downregulated genes in OM mainly belonged to both cellular and metabolic process in GO biological process gene sets.

As the embryo size was highly variable within each group, a PCA analysis was made on DEG in ICM and TE to verify that observed differences were effectively due to maternal age more than to embryo size (Figure 2b). Respectively 3 YM and 3 OM were classified in small (<1100µm in diameter) while the remaining 2 YM and 3 OM were classified in large group (> 1100µm in diameter).

In the ICM, the first axis of the PCA explained 58.3% of the variance of the data and clearly separated OM from YM, indicating than more than half of the variance in DEG is due to maternal aging. The second axis slightly separated genes according to embryo size but explained only 11.9% of the variance, indicating that in the ICM, embryo size explains a minor part of the variance. These observations were confirmed by the Partial redundancy analysis. Indeed, in ICM, 43% of the variance was explained by maternal age while only 14% was explained by embryo size. Moreover, the 20 variables that contributed the most to the separation are mostly explaining differences related to maternal age.

In the TE part, both the first and the second axis (explaining respectively, 67.3% and 16.5% of data variability) separated both maternal age and embryo size groups. The partial redundancy analysis also showed a less clear impact of maternal age in DEG from TE. Indeed, mares’ age explained 31% of the variance of DEGs whereas 22% was explained by embryo size. Moreover, although most of genes showed a clear contribution in the construction of the first axis, some such as cystathionine beta-synthase (*CBS*) and CD69 molecule (*CD69*) were clearly involved in the construction of the second axis.

After Entrez Gene ID conversion, 12,344 genes were considered expressed in ICM. In the GO Biological Process database, 106 identified gene sets differed according to maternal age in the ICM (11 enriched in OM and 95 enriched in YM) (Supplementary Table S5). In the KEGG database, 14 gene sets were perturbed (Supplementary Table S5).

Using SUMER analysis (Figure 5), gene sets enriched in OM ICM were mainly grouped under terms “Respiratory electron transport chain” and “Translational elongation”. The first term represents mitochondrial function while the second represents translation function. Moreover, terms relative to “Systemic lupus erythematosus” are conserved by SUMER analysis and enriched in OM ICM. Genes involved in these enrichments were also present in dysregulated metabolic pathways.

**Figure 5:**
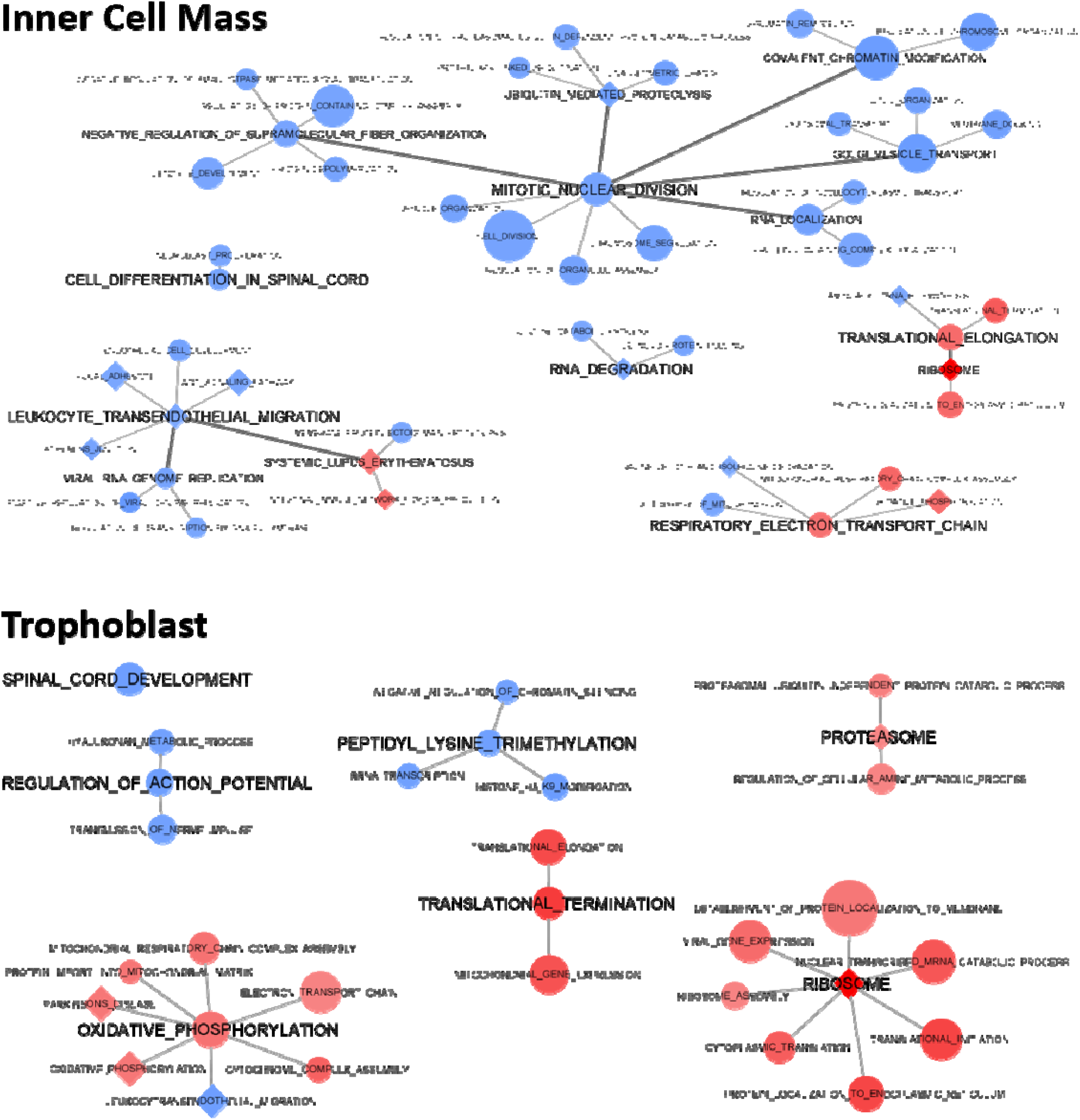
GSEA clustering of the most perturbed terms in ICM and TE according to mares’ age Nodes represent altered gene sets in ICM and TE (FDR < 0.05). Node size represents the gene set size. Node shape represents the gene set database: GO BP (circle) or KEGG (diamond). Blue nodes represent enriched gene sets in YM (NES < 0) while red nodes represent enriched gene sets in OM (NES > 0). Edges represent the level of connection between representative gene sets. This graph was performed using SUMER R package ^114^ and modified using cytoscape 3.8.2 ^115^ ICM: Inner Cell Mass; TE: trophoblast; FDR: False Discovery Rate; GO BP: Gene Ontology Biological Process; Kyoto Encyclopedia of Genes and Genomes; NES: Normalized Enrichment Score; YM: Young mares; OM: Old mares

Gene sets enriched in YM ICM were involved in different biological processes such as “Mitotic nuclear division” which is relative to mitosis. Under this term, “Spindle organization” and “Negative regulation of supramolecular fiber organization” terms represent the assembly of spindle during mitosis. “Chromosome segregation” is also found in this grouping. Furthermore, terms under “Golgi vesicle transport” seem to be involved in cytokinesis and vesicle trafficking. The catabolism of protein and mRNA is represented by “Ubiquitin mediated proteolysis”. Translation function is represented by several terms grouped under “covalent chromatin modification” in the mitosis part.

“Leukocyte transendothelial migration” term is also enriched in YM ICM and seems to represent cell signaling and adhesion. Indeed, this term regroups several pathways such as “Focal adhesion”, “Adherens junction” and “WNT signaling pathway”.

Finally, further terms related to embryo development could be found in different groups such as “Dendrite development”, “Cell differentiation in spinal cord” or “endothelial cell development”.

After Entrez Gene ID conversion, 11,874 genes were considered expressed in TE from OM or YM embryos. In GO BP database, 38 gene sets were perturbed by mares’ age in the TE of embryos (11 enriched in YM and 27 enriched in OM) (Supplementary Table S6). In the KEGG database, 5 pathways were enriched in the TE of OM embryos (Supplementary Table S6).

As in ICM part, using SUMER analysis, translation (represented by both “Translational termination” and “Ribosome” terms) and mitochondrial function (represented by “Oxidative phosphorylation” associated terms) were enriched in OM embryos. “Proteasome” term is enriched in OM embryos representing protein catabolism.

Moreover, transcription (represented by “Peptidyl lysine trimethylation” term) and embryo development (represented by “spinal cord development”) were, as in ICM, also, enriched in YM embryos. Finally, in TE, “Regulation of action potential” term and associated terms were enriched in YM. These terms could be related to ion movement.

## Discussion

Maternal aging did not affect embryo recovery rate (ERR), embryo size nor total RNA quantity. It is important to notice that, as the experimental protocol required only 5 embryos per group, a limited number of embryo collection attempts were performed, resulting in a limited statistical power for these analyses. ERR, especially in young mares, was not as good as in commercial embryo transfer programs ^31,36^. Moreover, total RNA content in equine embryo was exponentially correlated to embryo diameter and was not affected by maternal age.

As ovulation check was only performed 48 hours after induction of ovulation, there is a bias in the estimation of the embryo age in the present study. This certainly partly explains the large variation in embryo size observed here. Different studies, however, have shown that embryo size is highly variable ^37–39^, even when ovulation determination is accurate to the nearest 6 hours ^40^. Accurately determining fertilization time *in vivo* is not possible and the size of 142 equine embryos recovered at 8 days post-ovulation was reported to range from 120 to 3,980µm with an average of 1,132µm ^30^. In the present project, all embryos remain within these limits.

Although embryo size could be considered as normal for Day 8 embryos, embryo sizes were highly variable and not equally distributed between groups. Embryo size was considered as a cofactor in the differential analysis. In addition, partial redundancy analysis showed that embryo size explained less than 25% of the variability of DEGs in both ICM and TE. As effects of maternal age on equine embryo growth are not so clear ^24–28^, an interaction between both effects on embryo gene expression can not be excluded. Maternal age, however, explained 43 and 31% of the total variability of DEGs in both ICM and TE, respectively, suggesting that identified DEGs were mainly due to maternal age effects. It is important to note that in ICM, the proportion of variability explained by maternal age was 3 times greater than that of embryo size, whereas in the TE, this difference was smaller. This could be explained by the fewer number of DEGs identified in TE compared to ICM but also because trophoblast expansion was possibly more correlated to embryo size than the ICM.

As in tumors, today, the only way to analyze separate expression of ICM and TE cells in mammalian embryos is to use micromanipulation but micromanipulation is expensive in time and requires adapted skills and material. In this study, micromanipulation to properly separate ICM and TE was not possible. Therefore, deconvolution appeared to be a good option to analyze the effects of maternal ICM gene expression in mixed ICM and TE samples. Deconvolution seemed to have been effective on embryo to separate ICM and TE gene expression. Indeed, the 341 genes differentially expressed in ICMandTE *vs* TE_part were also identified by comparing TE with post-deconvolution ICM cells gene expression dataset. Moreover, after deconvolution, more genes were differentially expressed between TE and ICM datasets and among them, more genes identified in literature were differentially expressed. Finally, PCA showed that deconvolution separate DeMixT_ICM_cells from other datasets while before deconvolution ICMandTE and TE_part were partly superposed. Only POU5F1 and UTF1 differed in terms of allocation between previously published data ^35^ and the present study. In data before deconvolution, expression of POU5F1 and UTF1 were extremely variable according to embryo diameter in both compartments. Deconvolution reduced the variability of both gene expression in ICM but not in TE. Studies showed that POU5F1 is expressed in both compartments until Day 10 post ovulation in equine in vivo produced embryos ^41^ but its expression decreases progressively from Day 7 in TE cells. Thus, the statistically increased expression of POU5F1 in TE on Day 8 is compatible with previously published data.

Therefore, it was chosen to only present the analysis of the effects of age in DeMixT_ICM_cells dataset. In all cases, deconvolution did not change the principal results of maternal aging on embryo gene expression because the comparison of maternal age in ICMandTE and DeMixT_ICM_cells gave comparable results.

In both ICM and TE, the expression of genes involved in mitochondrial function and translation with ribosome biogenesis were enriched in OM embryos while transcription with chromatin modification was enriched in YM embryos (Figure 6). Mitosis, signaling and adhesion pathways were additionally altered and the expression of genes involved in embryo development was particularly reduced in the ICM hemi-embryos from old mares. Finally, in the TE, ion movement related genes were affected. Several studies on mature oocytes in women and mice, using microarrays and RNAseq, had shown a differential expression of genes involved in mitochondrial, cell cycle, cytoskeleton functions and related pathways according to maternal age ^42–44^, as observed in the present study in equine blastocysts, especially in the ICM. In equine follicles and oocytes, the expression of genes essential for oocyte maturation, as studied by RT-qPCR, quantitatively and temporally differed with maternal age, suggesting a desynchronization ^16^. Moreover, a study on the expression of 48 genes that have been previously identified as related to aneuploidy and maternal aging in humans showed differential expression in cumulus-oocytes complexes from young (< 12 years of age) and old (> 18 years old) mares ^17^. Genes related to mitochondrial replication and function were also down-regulated in oocytes of old mares ^23^. Even effects of maternal aging on gene expression in equine oocytes, using high throughput techniques, are not yet available, results on equine Day 8 blastocysts suggest that mitochondrial perturbations present in the oocyte persist until the blastocyst stage. In humans, single-embryo RNA-seq analysis showed that maternal aging alters the transcriptome of blastocysts derived from ICSI. Genes altered by maternal age were related to segregation of chromosomes, embryonic growth and cell signaling ^45,46^, as observed in our study on equine blastocysts.

**Figure 6:**
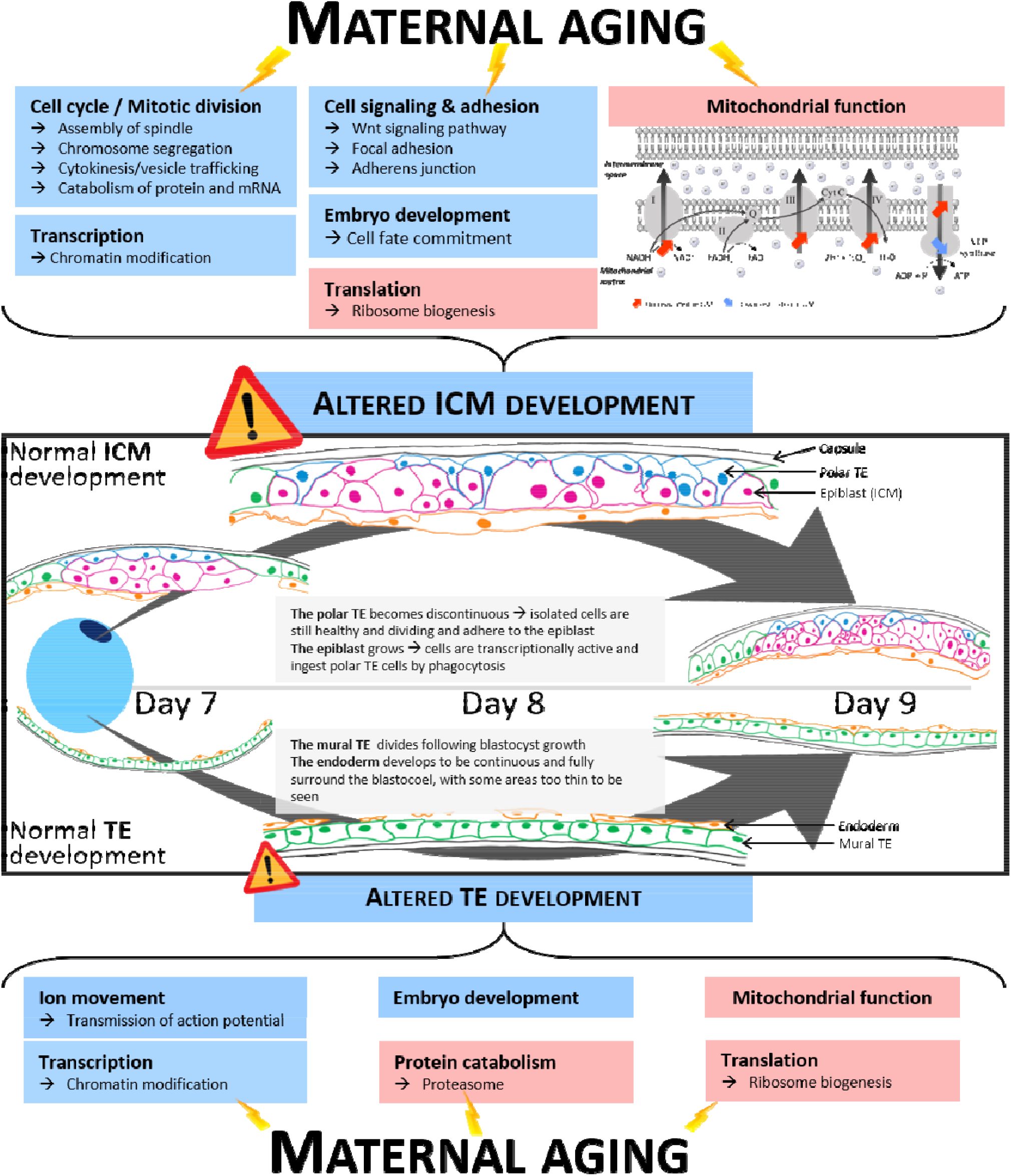
Schematic representation of the observed effects of maternal aging on ICM and TE in equine blastocysts Normal blastocyst development from day 7 to day 9 is represented inside black square using literature ^47,48^. Light blue boxes represent biological processes that are enriched in YM embryos and light red boxes represent biological processes that are enriched in OM embryos.

Unlike the human embryo, the equine embryo remains spherical, grows very fast and is surrounded by its glycoprotein capsule up to 22 days post ovulation ^37^. At 8 days post ovulation, equine embryos are therefore growing and the endodermal layer entirely delineates the future yolk sac (Figure 6) ^47^. Moreover, whereas in Day 7 embryos, the epiblast is lined by a continuous layer of trophoblast named the polar trophoblast. As the blastocyst grows, epiblast cells phagocyte polar trophoblast cells that become discontinuous. During this time, epiblast and polar trophoblast cells divided and are bound by tight junctions ^48^. The turnover in focal adhesion is essential for cell movement and phagocytosis ^49^.

Cell signaling and adhesion function were altered in OM embryos. Indeed, KEGG pathways “focal adhesion” and “adherens junction” regrouped in “Leukocyte transendothelial migration” on Figure 4 are disturbed in old mares’ embryos. Focal adhesions are subcellular structures observed in adherent cells, that are clusters of different structural and signaling molecules and integrins that form a physical link between the actin cytoskeleton and the extracellular matrix ^50,51^. Integrins are the major receptors for cell adherence to extracellular matrix proteins in focal adhesions by playing an important role in transmembrane connections to the cytoskeleton and activation of several signaling pathways, as growth factor receptors ^52^. Integrin Subunit Alpha V (*ITGAV*) expression was reduced in the ICM of embryos collected from OM. In different human breast cancer cell lines, cell proliferation, invasion and renewal are restricted when *ITGAV* is inhibited ^53^.

Moreover, the activity and localization of small GTPases within the Rho family must be precisely regulated as the communication between them is indispensable to achieve the turnover of actin filaments and cell adhesions necessary for cell migration, that are essential processes for the proper development of the embryo ^54^. The Rho GTPase Activating Protein 9 and 35 (*ARHGAP9* and *ARHGAP35*) were downregulated in OM ICM. *ARHGAP35* codes for P190 Rho GTPase activating protein that mediates signalization through integrin-dependent adhesion in cells. Aberrant neural morphogenesis, related to an accumulation of polymerized actin in embryoneural tube cells, is reported in mice lacking P190 Rho GTPase activating proteins ^55^. Furthermore, P21 activated kinases (PAK) are effector kinases for other small Rho GTPases such as Rac that regulate the polymerization of actin and can affect the microtubule organization ^56^. Once activated, PAKs promote the turnover of focal adhesions by regulating transcription ^57^. *PAK3* was downregulated in the OM ICM. However, in TE, SLIT-ROBO Rho GTPase Activating Protein 3, *SRGAP3* was one of the upregulated genes in OM. This gene regulates cytoskeletal reorganization and plays an important role in the formation of cell-cell adhesion ^58^. Therefore, maternal aging seems to alter focal adhesion in both ICM and TE of embryos but, while in ICM, it seems to reduce cohesion and signalization, in TE it seems to reinforce them.

Several signaling pathways required for development were altered and numerous genes related to focal adhesions and signaling pathways were down-regulated in OM ICM (*ARHGAP9* and *ARHGAP35*; B-Raf Proto-Oncogene, Serine/Threonine Kinase, *BRAF*; Filamin C, *FLNC*; FYN Proto-Oncogene, Src Family Tyrosine Kinase, *FYN*; *PAK3*; Protein Phosphatase 1 Regulatory Subunit 12A, *PPP1R12A*; Talin 2, *TLN2*; X-Linked Inhibitor Of Apoptosis, *XIAP*). Among all signaling pathways, transforming growth factor beta (TGF-β), Wg et Int (Wnt), mitogen-activated protein kinases (MAPK) and apoptosis signaling were altered by maternal age in equine embryos. Indeed, transforming growth factor beta receptor 1 (*TGFBR1*) was downregulated in the ICM of embryos from old mares. *TGFBR1* is one of the two dimers for the receptor for transforming growth factors beta that is essential in the generation of axes and cell fate commitment during mammal embryogenesis ^59^. Moreover, Casein Kinase 1 Alpha 1, (*CSNK1A1*) and *Protein* Phosphatase 2 Regulatory Subunit B’Epsilon (*PPP2R5E*), involved in Wnt signaling pathway, were downregulated in OM embryos. Wnt signaling pathway is also involved in nervous development system and may act on cell determination in mice embryos ^60^. Moreover, Doublesex And Mab-3 Related Transcription Factor 3 (*DMRT3*) was downregulated in the OM ICM. *DMRT3* is required for the proper development of the brain in prenatal chicken and mice ^61^.

Cell cycle related pathways and genes were also disturbed by maternal aging. Indeed, the expression of Pleckstrin Homology domain-Interacting Protein (*PHIP*), also known as replication initiation determinant, was reduced in the OM ICM. This gene is required for the initiation of DNA replication. Fibroblasts originated from mice embryos with depleted *PHIP* expression presented less replication-initiation as well as abnormal replication fork progression events ^62^. Moreover, several cyclins (CCN) were less expressed in the ICM of embryos from old mares (*CCNB3, CCNI* and *CCNT2*). Cyclins are key cell cycle regulators and, for instance cyclin B3 degradation allows the passage from metaphase (chromatids aligned on metaphase plate) to anaphase (segregation of sister chromatids) ^63^. During metaphase and anaphase, spindle assembly and chromosome segregation could also be compromised. Indeed, genes involved in the formation of microtubules (Tubulin Gamma 2, *TUBG2*) ^64^, the generation of the mitotic spindle (Never in Mitosis A related kinase 9, *NEK9*) ^65^, the production of cohesion and condensing complexes (Structural Maintenance Of Chromosomes 4, *SMC4*) ^66^, the control of centriole length, important for centrosomes and microtubules organization (Centriolar Coiled-Coil Protein 110, *CCP110*) ^67^, the amplification of intra-spindle microtubules allowing the formation of a sufficient quantity of kinetochore (HAUS Augmin Like Complex Subunit 6, *HAUS6*) ^68^, the stabilization and protection of the cohesin complex association with chromatin (Cell Division Cycle Associated 5, *CDCA5*; Shugosin 2, *SGO2*) ^69,70^ and the production of microtubule-based motor proteins (Kinesin Family Member 14 and 23, *KIF14* and *KIF23*) ^71^ were downregulated in OM ICM. Furthermore, Centromere protein F (*CENPF*), an important regulator of chromosome segregation during mitosis, was reduced in OM ICM. CENPF is indispensable for murine embryo development from the first stages of development until at least the blastocyst stage ^72^. Impaired *CENPF* expression could induce chromosome mis-segregation in the ICM of old mares’ embryos.

Cytokinesis seems to also be altered by maternal age. CD2-associated protein (*CD2AP*) and Vacuolar Protein Sorting 4 Homolog A (*VPS4A*) were downregulated in OM ICM. Both are involved in the late phase of cytokinesis which correspond to the abscission of the cell, producing two daughter cells ^73,74^. Globally, results suggest that cell division is impaired or indicate that there is less division in the ICM of old mares’ embryos.

Transcription is regulated by epigenetic marks. Several pathways linked to methylation and acetylation were enriched in YM ICM and TE in relation to numerous upregulated genes in YM ICM (i.e., Histone Deacetylase 1 and 2, *HDAC1* and *HDAC2*; Helicase, Lymphoid Specific, *HELLS*; DNA Methyltransferase 3 Alpha, *DNMT3A*; Lysine Acetyltransferase 6B, *KAT6B*; Lysine Demethylase 4A, *KDM4A*; Lysine Methyltransferase 2A, *KMT2A*; Nuclear Receptor Binding SET Domain Protein 3, *NSD3*; SET Domain Containing 5, *SETD5*; Tet Methylcytosine Dioxygenase 1, *TET1*). At blastocyst stage, *de novo* methylation is more important in the inner cell mass than in trophoblast cells and is essential for the proper development of mammalian embryos ^75^. *DNMT3A* is essential for *de novo* methylation and required for embryo viability ^76^. As it has been shown that maternal aging increases the acetylation of lysine 12 of histone H4 in mice oocytes, affecting fertilization and embryo development ^77^, it is likely that the methylation and acetylation dysregulation observed in the embryos of old mares could be inherited from the oocyte rather than directly due to embryo environment.

Both in OM ICM and TE, several pathways related to mitochondrial function were enriched. Mitochondria are long known to be the “powerhouse” of cells, producing most of the necessary ATP for cellular function ^78^. ATP production is possible through oxidative phosphorylation enabled by the electron transport chain in the inner membrane of mitochondria. This process begins with the oxidation of NADH or FADH that generates a proton gradient across the mitochondrial membrane, driving ATP synthesis through the phosphorylation of ADP *via* ATPase ^79^. Enriched pathways in OM embryos demonstrate that the NADH dehydrogenase (complex I), the oxidoreductase acting on a heme group of donors (complex III) with the global electron carrier, and the proton-transporting ATP synthase (1^rst^ part of the complex V) activities were particularly enriched while the second part of complex V is depleted. Since all complexes are required to produce ATP, the impaired complex generation may lead to mitochondrial stress and uncoupled respiratory chain, resulting in poor ATP production. In human IVF embryos, reduced mitochondrial respiration efficiency, with a negative effect on embryo development, is observed when oocytes are collected from old women ^80^. In mammalian expanded blastocysts, energy requirement is high, probably because of the need for ion transport enabling quick blastocoel cavity expansion. Defects in metabolic requirements during blastocyst development are associated with impaired implantation ^81^. Equine embryos are not an exception ^82^.

Mitochondria being maternally inherited, uncoupling of the mitochondrial respiratory chain may originate from mitochondrial defects already present in oocytes ^83^. Studies have shown that, in human, poor oocyte quality is responsible for the decline in fertility associated with maternal aging. The reduced oocyte developmental competence in older women is related to mitochondrial dysfunction ^84^. In horses’ oocytes, before *in vitro* maturation, no difference between young and old mares is observed while after, reduced mitochondrial DNA copy numbers, increased swelling and reduced mitochondrial cristae are reported in oocytes collected from mares >12 years. This suggests that oocyte mitochondria from old mares present deficiencies that become problematic when energy needs are high, as during *in vitro* maturation or subsequent embryo development ^21^. Because mitochondrial replication in equine embryos does not begin until the blastocyst stage ^85^, oocyte mitochondrial numerical or functional deficiencies may compromise early embryogenesis.

In TE, the GO biological process “Regulation of action potential” and “Transmission nerve impulse”, related to ion movement, were enriched in YM embryos. These processes are associated with genes involved in the action potential dependent ion channels. These ion channels such as NA^+^, K^+^, ATPase are essential for the accumulation of blastocoel and subsequently yolk sac fluid in equine embryos ^86^. Their inhibition is detrimental for embryo development ^87^. Thus, both reduced ATP production and disturbed ion channels’ function may impact embryo growth, although effects of mares’ age on embryo growth are controversial ^24–28^.

This is the first study showing an effect of maternal age on gene expression in the equine blastocyst at Day 8 post ovulation. Maternal age, even for mares as young as 10 years old, disturbs several processes related to chromatin segregation, cytokinesis, apposition of epigenetic marks and mitochondrial function and that the ICM appears to be more affected than the TE. These perturbations on Day 8 post ovulation may affect the further development of the embryo and as a result, may contribute to decreased fertility due to aging. So far, maternal age does not appear to have an impact on foal morphology at birth but, as for placental function, more subtle effects may exist. Here, as embryos were collected long before implantation, it is impossible to know if these embryos would have implanted and if they would have produced a live foal or not. Studies on long-term growth and health of foals born to the same mares are currently being performed.

## Supporting information

Supplementary infos

## Acknowledgments

The authors are grateful to the staff of the Institut Français du Cheval et de l’Equitation (IFCE) experimental farm (Domaine de la Valade, Chamberet, France) for care and management of animals. We acknowledge the High-throughput sequencing facility of I2BC for its sequencing and bioinformatics expertise. The bioinformatics analyses were performed thanks to Core Cluster of the Institut Français de Bioinformatique (IFB) (ANR-11-INBS-0013). Many thanks to Matthias Zytnicki and Christophe Klopp for their advice on RNA-seq de novo analysis. Many thanks to Pablo Ross who kindly provided the coordinates for the XIST gene. The authors thank Denis Laloe for its help with vegan package. The authors thank Shavahn Loux for her information on PLAC8A. The authors are grateful to Alice Jouneau, Sophie Calderari and Patrice Humblot who kindly revised this article.

## Author contributions

PCP obtained the funding. PCP and VD conceived the project. LW, PCP, and VD supervised the study. ED, CD, CA, ND, NP, LW, VD and PCP adapted the methodology for the project. ED, CD, CA, YJ, JAR, and LW performed the experiments. CD, CA, ND, NP, MD and LW provided the resources. ED, LJ, YJ and RL performed data curation. ED and LJ analysed the data. ED wrote the original draft. All authors read, revised, and approved the submitted manuscript.

