## Supplementary figures and images for "Maternal age affects equine Day 8 embryo gene expression both in trophoblast and inner cell mass"

### Additional file 1

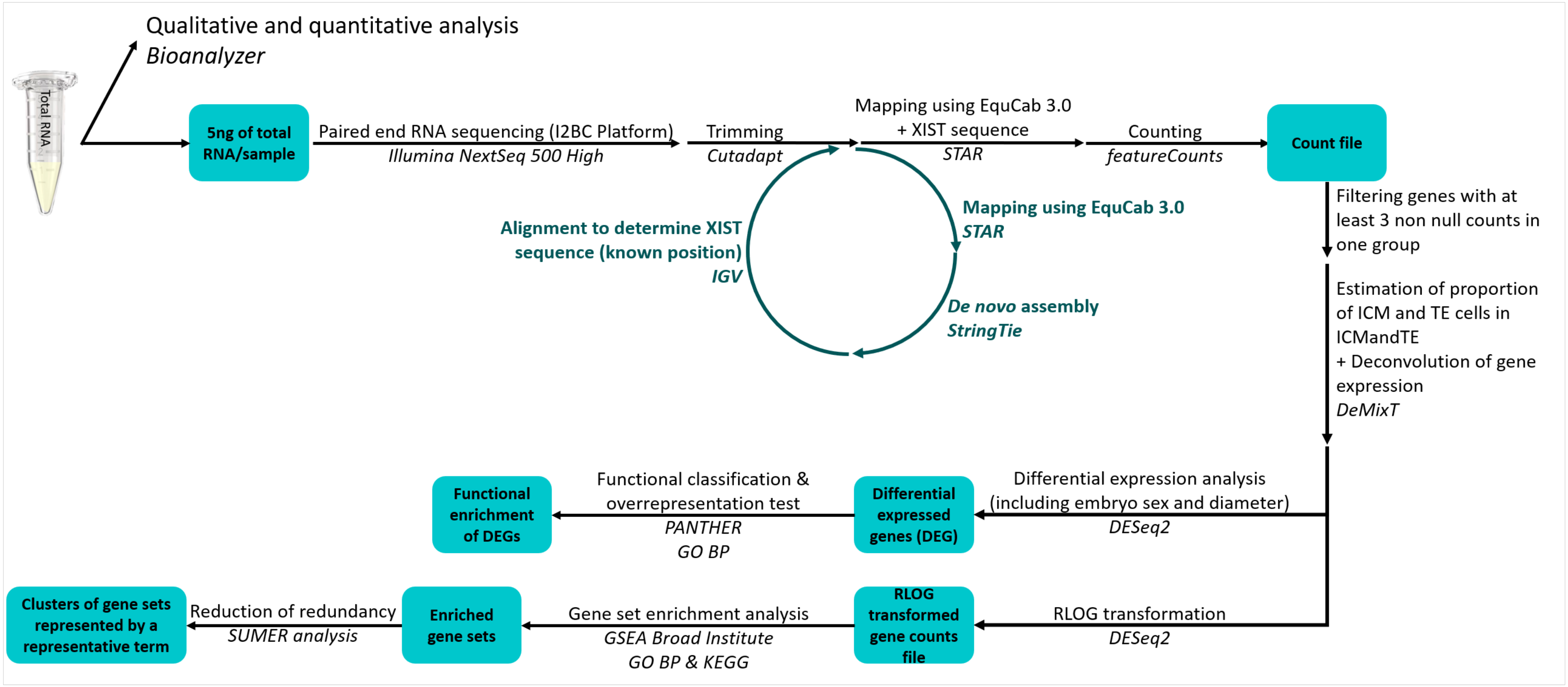
