## Additional file 2 for "Maternal age affects equine Day 8 embryo gene expression both in trophoblast and inner cell mass"

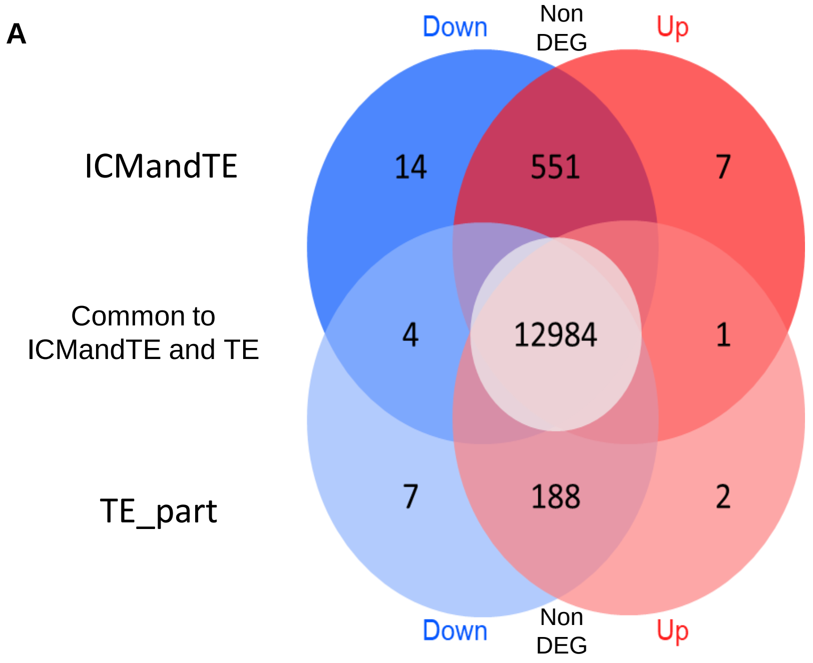


**B**

| | ICM and TE | pure TE |
|---|---|---|
| CST6 | 0.96 | 0.87 |
| ENSECAG00000035582 | 0.8 | 0.75 |
| GALM | 0.66 | 0.61 |
| GTPBP8 | 0.69 | 0.58 |
| GNG10 | 0.42 | 0.37 |
| NMI | 2.28 | 0.35 |
| MGST1 | 0.47 | 0.35 |
| USP11 | -0.53 | -0.15 |
| PARP1 | -0.56 | -0.43 |
| DESI1 | -0.57 | -0.3 |
| MARS2 | -0.59 | -0.14 |
| MKI67 | -0.59 | -0.42 |
| MYH10 | -0.62 | -0.35 |
| PCM1 | -0.64 | -0.28 |
| PPFIBP1 | -0.72 | -0.05 |
| H1-4 | -0.79 | -0.87 |
| PHKA2 | -1.06 | -0.6 |
| KMT2A | -1.1 | -0.17 |
| MAP1B | -1.41 | 0.42 |
| DPYSL5 | -2.12 | -0.53 |
| PAK3 | -3.91 | |

**C**

| | ICM and TE | pure TE |
|---|---|---|
| CD69 | | 5.09 |
| PFAS | -0.34 | -0.38 |
| CBS | -0.49 | -0.37 |
| SRGAP3 | -0.55 | 2.52 |
| SETX | -0.56 | -0.49 |
| LUZP1 | -0.72 | -0.75 |
| REV3L | -0.37 | -0.75 |
| ATAD2B | -0.52 | -0.93 |
| ENSECAG00000022280 | -1.7 | -2.53 |

**D**

| | ICM and TE | pure TE |
|---|---|---|
| RPS29 | 0.95 | 0.78 |
| AASS | -0.89 | -0.7 |
| H4C3 | -1.15 | -1.33 |
| RNASE1 | -2.06 | -2.14 |
| ENSECAG00000038654 | -1.96 | -2.48 |

log2FC -2.5 0.0 2.5 5.0

**<u>Venn diagram (A) and tables of DEG in TE_part and ICMandTE according to maternal age (B, C and D)</u>**

Table B presents DEGs in ICMandTE, Table C presents DEGS in TE_part and Table D presents DEGS in common in both parts. Blue represents down regulated DEG (log2FC<0) while red show up regulated DEG (log2FC>0) in OM embryos compared to YM embryos. DEGs are in Entrez Gene ID or Ensembl Equine Gene ID (ENSECAG) when Entrez Gene ID was not available.

DEG: Differentially Expressed Genes (padj<0.05); log2FC: Fold change in log2; TE: Trophoblast; ICM: Inner Cell Mass; OM: group of embryos produced by Old Multiparous mares; YM: group of embryos produced by Young Multiparous mares

### ICMandTE

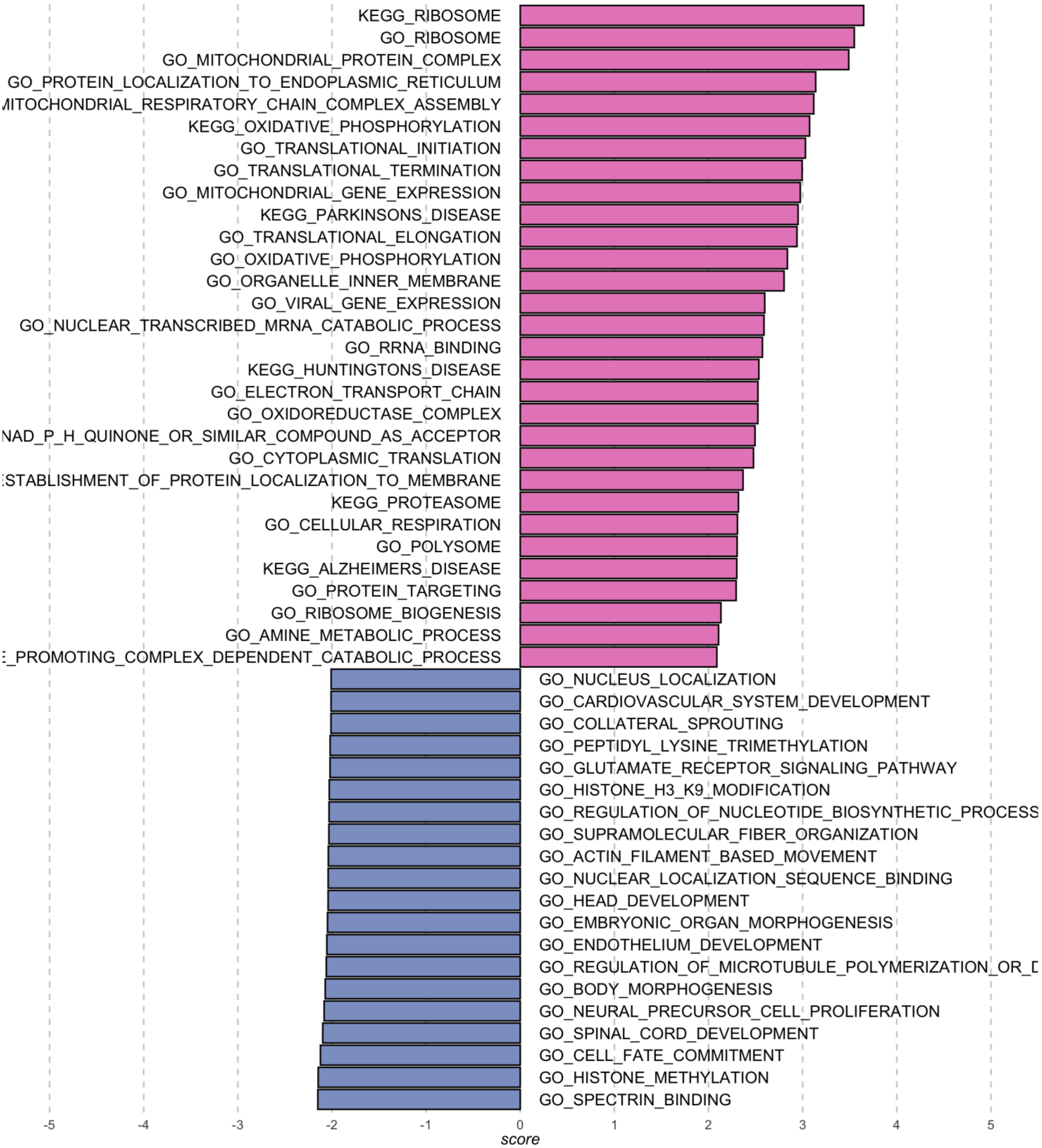


**<u>Diagram of most perturbed GO and KEGG pathways (FDR < 0.05) in inner cell mass enriched part (ICMandTE) of equine D8 blastocysts according to maternal age</u>**

Pink pathways are the most enriched pathways in embryos from old multiparous mares and purple pathways are the most enriched pathways in embryos from young multiparous mares. Score represents False Discovery Rate using a Gene Set Enrichment Analysis.

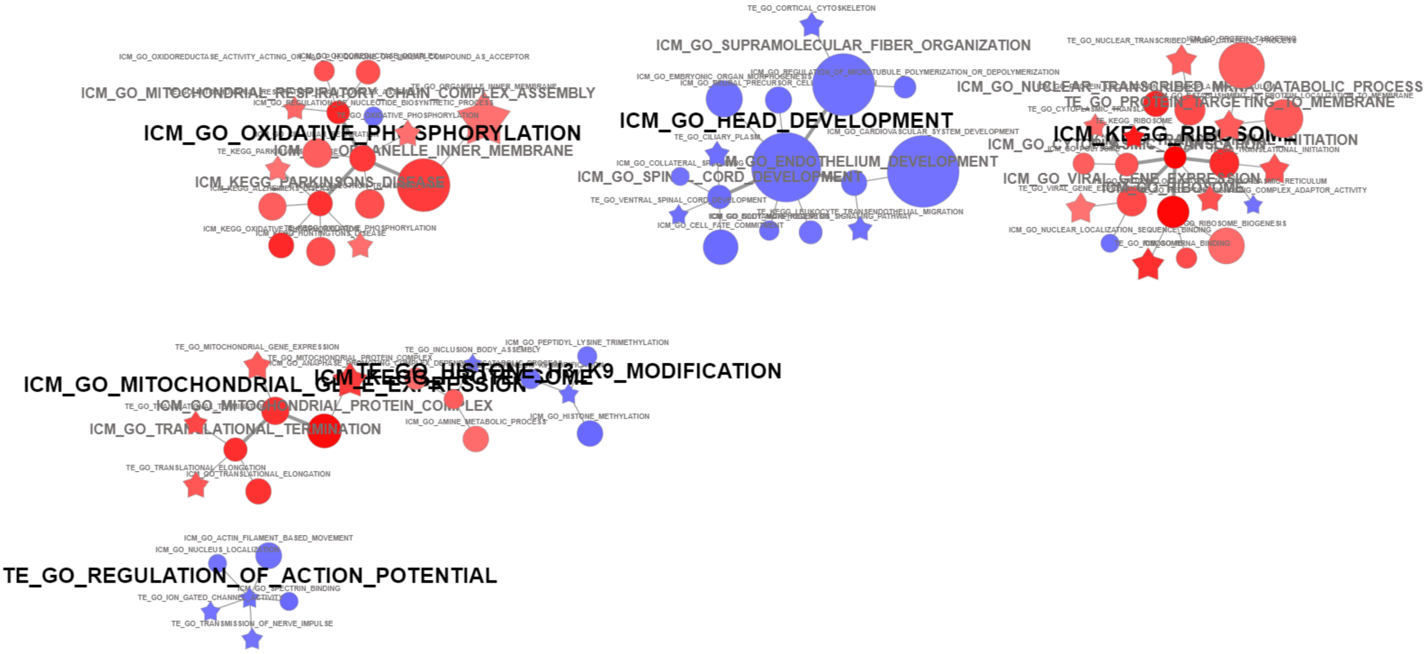


**<u>Clustering of the most perturbed GO and KEGG terms in ICMandTE and TE  according to mares' age</u>**

Graph representing in nodes GO Biological process and KEGG representative terms in ICMandTE (circle) and TE_part (star). Node size represents the size of the gene set. Blue nodes represented gene sets enriched in YM (NES < 0) while red nodes represented gene sets enriched in OM (NES > 0). Edges represent the level of connection between representative gene sets. This graph was performed using SUMER R package  [55].

NES: Normalized Enrichment Score; ICM: Inner Cell Mass; OM: group of embryos produced by Old Multiparous mares; YM: group of embryos produced by Young Multiparous mares
