## Supplementary infos for "Maternal age affects equine Day 8 embryo gene expression both in trophoblast and inner cell mass"

**Supplementary information**

Supplementary Figure S1:

Supplementary Figure 1.png

Pipeline of biostatistical analysis

Once extracted, total RNA qualification and quantification were obtained using a Bioanalyzer. For sequencing, 5 ng/sample of total RNA were used for paired end RNA sequencing (I2BC platform) using Illumina NextSeq 500 high technology. Sequences were trimmed using Cutadapt. To determine XIST sequence, a *de novo* assembly was made using a mapping with STAR and StringTie. The GTF observed was aligned in IGV and XIST sequence was found at the known position. XIST sequence was included in EquCab 3.0 and a new mapping was performed using STAR. Counting was performed using featureCounts. The differential analysis was performed including embryo sex and diameter using Deseq2. A functional enrichment analysis of DEGs was performed using PANTHER web software. In another time, counts of all genes were normalized using Deseq2 and a gene set analysis (GSEA) was performed using the Gene Ontology (GO) biological process (BP) and the Kyoto Encyclopedia of Genes and Genomes (KEGG) databases with GSEA software form the Broad Institute. To reduce redundancy between terms, SUMER analysis was performed on enriched gene sets.

Supplementary Figure S2:

Supplementary Figure 2.pdf

Analysis of ICM enriched and TE part before gene expression deconvolution using DeMixT

On first page, the analysis of differential expressed genes is represented. The second page represent the results of the gene set enrichment analysis of ICMandTE gene expression. The third page presents the clustering of gene sets altered by maternal age from SUMER analysis of the GO BP and KEGG terms of mixed ICMandTE and TE_part.

Supplementary Table S1:

Supplementary Table 1.csv

Differential gene analysis using DeSeq2 in ICMandTE of equine embryo at Day 8 post-ovulation

Equine ensemble ID, orthologue human Ensembl ID, Orthologue human Entrez Gene ID, gene description, normalized counts for each sample of ICM and parameters obtained after Deseq2 analysis (log2FoldChange, pvalue and padj (after FDR correction)) of genes expressed in ICM enriched part (before gene expression deconvolution using DeMixT) of OM and YM embryos

ICM: Inner cell mass; OM: old mares; YM: young mares

Supplementary Table S2:

Supplementary Table 2.csv

Gene set enrichment analysis results on gene expression comparing ICMandTE of embryos from old mares to young mares

Gene Set Enrichment Analysis results (pathway name, GO accession when possible and size, Normalized Enrichment Score, p-value and FDR corrected q-value) for GO biological process and KEGG databases on ICMandTE gene expression table.

Supplementary Table S3:

Supplementary Table 3.csv

Differential gene analysis using DeSeq2 in DeMixT_ICM_cells of equine embryo at Day 8 post-ovulation

Equine ensemble ID, orthologue human Ensembl ID, Orthologue human Entrez Gene ID, gene description, normalized counts for each sample of ICM and parameters obtained after Deseq2 analysis (log2FoldChange, pvalue and padj (after FDR correction)) of genes expressed in ICM (after gene expression deconvolution of ICMandTE using DeMixT) of OM and YM embryos

ICM: Inner cell mass; OM: old mares; YM: young mares

Supplementary Table S4:

Supplementary Table 4.csv

Differential gene analysis using DeSeq2 in TE_part of equine embryo at Day 8 post-ovulation

Equine ensemble ID, orthologue human Ensembl ID, Orthologue human Entrez Gene ID, gene description, normalized counts for each sample of ICM and parameters obtained after Deseq2 analysis (log2FoldChange, pvalue and padj (after FDR correction)) of genes expressed in TE of OM and YM embryos

TE: Trophoblast; OM: old mares; YM: young mares

Supplementary Table S5:

Supplementary Table 5.csv

Gene set enrichment analysis results on gene expression of DeMixT_ICM_cells of embryos from young and old mares

Gene Set Enrichment Analysis results (pathway name, GO accession when possible and size, Normalized Enrichment Score, p-value and FDR corrected q-value) for GO biological process and KEGG databases on DeMixT_ICM_cells gene expression table (after gene expression deconvolution on ICMandTE using DeMixT).

Supplementary Table S6:

Supplementary Table 6.csv

Gene set enrichment analysis results on gene expression of TE_part of embryos from young and old mares

Gene Set Enrichment Analysis results (pathway name, GO accession when possible and size, Normalized Enrichment Score, p-value and FDR corrected q-value) for GO biological process and KEGG databases on TE_part gene expression table.
